# Efficient SARS-CoV-2 detection utilizing chitin-immobilized nanobodies synthesized in *Ustilago maydis*

**DOI:** 10.1101/2022.11.11.516239

**Authors:** Magnus Philipp, Lisa Müller, Marcel Andrée, Kai P. Hussnaetter, Heiner Schaal, Michael Feldbrügge, Kerstin Schipper

**Affiliations:** Institute for Microbiology, Heinrich Heine University Düsseldorf, Universitätsstraße 1, 40225 Düsseldorf, Germany; Institute of Virology, Medical Faculty, Heinrich Heine University Düsseldorf, Universitätsstraße 1, 40225 Düsseldorf, Germany

**Author notes:** Corresponding author: Kerstin Schipper.

**Keywords:** Nanobody, SARS-CoV-2, chitin, chitinase, *Ustilago maydis*

## Abstract

The COVID-19 pandemic has greatly impacted the global economy and health care systems, illustrating the urgent need for timely and inexpensive responses to a pandemic threat in the form of vaccines and antigen tests. The causative agent of COVID-19 is SARS-CoV-2. The spike protein on the virus surface interacts with the human angiotensin-converting enzyme (ACE2) via the so-called receptor binding domain (RBD), facilitating virus entry. The RBD thus represents a prime target for vaccines, therapeutic antibodies, and antigen test systems. Currently, antigen testing is mostly conducted by qualitative flow chromatography or via quantitative ELISA-type assays. The latter mostly utilize materials like protein-adhesive polymers and gold or latex particles. Here we present an alternative ELISA approach using inexpensive materials and permitting quick detection based on components produced in the microbial model *Ustilago maydis*. In this fungus, heterologous proteins like biopharmaceuticals can be exported by fusion to unconventionally secreted chitinase Cts1. As a unique feature, the carrier chitinase binds to chitin allowing its additional use as a purification or immobilization tag. In this study, we produced different mono- and bivalent SARS-CoV-2 nanobodies directed against the viral RBD as Cts1 fusions and screened their RBD binding affinity *in vitro* and *in vivo*. Functional nanobody-Cts1 fusions were immobilized on chitin forming an RBD tethering surface. This provides a solid base for future development of an inexpensive antigen test utilizing unconventionally secreted nanobodies as RBD trap and a matching ubiquitous and biogenic surface for immobilization.

## 1 Introduction

The current COVID-19 pandemic challenges not only global healthcare systems and economies but has also underlined the strong demand for novel and versatile strategies to fight viral pandemics. In this regard, major innovations have already been driven by the pandemic, exemplified by the prompt development of mRNA-based vaccines (Kudlay and Svistunov 2022). Furthermore, the adaptation of monoclonal antibody therapeutics formerly mostly used in cancer patients for the treatment of COVID-19 represented an important step (Sun and Ho 2020, Bierle et al. 2021).

COVID-19 is caused by SARS-CoV-2. With the onset of the pandemic, the structure of the virus has been elucidated both on RNA (Jain et al. 2020) and protein level (Korber et al. 2020, Ou et al. 2020, Walls et al. 2020, Wrapp et al. 2020b). The spike protein complex was identified as a key player, as it is not only exposed on the surface of the viral particle, but also enables the attachment of the virus to the host cell via the human Angiotensin receptor 2 (ACE2) (Wang et al. 2020a). This mechanism has also been observed for other beta corona viruses like SARS-CoV (Hulswit et al. 2016). Spike proteins of these viral species usually consist of the two main subunits S2 and S1. S2 mainly serves as anchor of the protein in the viral membrane and also mediates fusion of the viral envelope and the host cell membrane (Hulswit et al. 2016). S1 is responsible for ACE2 binding (Wang et al. 2020a). Corona virus S1 proteins are generally organized into four domains, of which domains A and B form the receptor binding domain (RBD) which mediates ACE2 binding (Li et al. 2003, Wang et al. 2020a). The B subdomain of the RBD carries an extended loop that is highly variable among corona virus species and therefore also referred to as hypervariable region (Kirchdoerfer et al. 2016). All SARS-CoV-2 variants of concern that have been structurally elucidated to date (B.1.1.7 Alpha, B.1.351 Beta and B.1617 Delta and B.1.1.529 Omicron) carry mutations within the RBD domain that are assumed to play a role in infectivity and transmissibility of the virus (Baral et al. 2021, Torjesen 2021, VanBlargan et al. 2022). Therefore, the spike protein and especially its RBD domain are key targets for the development of therapeutics and vaccines.

The majority of vaccines cleared for use to date use an mRNA template of the spike protein that is translated in the host to evoke an immune response (Callaway 2020, Fernandes et al. 2022). However, since it was realized that vaccinated persons can still be infected with and spread SARS-CoV-2, there is a strong pressure to further develop test systems and therapeutics for a multi-layered strategy for COVID-19 treatment and control of SARS-CoV-2 spreading. Antibodies are key to both test systems and drug development. *Camelidae* and shark derived single heavy chain antibodies and derived nanobodies are emerging as potent alternatives to conventional antibodies (Muyldermans 2013, Salvador et al. 2019). *Camelidae* type antibodies only carry a heavy chain on their IgG scaffold as opposed to the light- and heavy chain of regular mammalian antibodies (Muyldermans 2013). This heavy chain alone (the so-called nanobody) can be quickly adapted to novel targets such as SARS-CoV-2 and production in microbial hosts is straightforward (Muyldermans et al. 2009, Wrapp et al. 2020a). Nanobodies have been shown to bind ligands in the nanomolar range and are stable under conditions of chemical and heat induced stress (Muyldermans 2013), which makes them promising molecules for widespread antigen testing. To this end several SARS-CoV-2 nanobodies engineered synthetically via phage display or generated directly by immunization of llamas, alpacas, and sharks have been published (Custodio et al. 2020, Gauhar et al. 2021, König et al. 2021).

We utilize the yeast form of the microbial model *Ustilago maydis* to produce heterologous proteins including alternative antibody formats like single chain variable fragments (scFvs) and nanobodies (Sarkari et al. 2014, Terfrüchte et al. 2017). Recently, we also established production of functional synthetic anti-SARS-CoV-2 nanobodies as a proof-of-principle for protein biopharmaceuticals (Philipp et al. 2021). For secretion of heterologous target proteins, a recently described unconventional secretion mechanism used by fungus to export chitinase Cts1 during cytokinesis is exploited (Reindl et al. 2019). Therefore, proteins of interest are fused to Cts1 which serves as a carrier for the export into the culture supernatant (Stock et al. 2012, Stock et al. 2016). Cts1 exhibits chitin binding activity making it a potential build-in immobilization- and purification tag (Terfrüchte et al. 2017). In addition, Jps1, a potential anchoring factor needed for Cts1 secretion is released into the culture medium and can be employed as alternative carrier (Philipp et al. 2021). Of note, proteins directed to the unconventional secretion pathway are not decorated with potentially harmful post translational protein modifications such as *N*-glycosylation which could lead to strong reactions in patients when proteins are applied as biopharmaceuticals (Stock et al. 2012).

Here, we exploited the dual functionality of chitinase Cts1 to produce different published SARS-CoV-2 nanobody versions via unconventional Cts1 secretion in *U. maydis*. Nanobody fusions were screened for their antigen binding activity *in vitro* and *in vivo*. Using the most promising binders, we established a novel strategy of RBD detection using a chitin surface for immobilization. In the future, these components can be combined to design a novel inexpensive and versatile virus detection system based on fungal compounds and a cognate biogenic chitin surface.

## 2 Results

### 2.1 Functional comparison of SARS-CoV-2 nanobody variants produced by unconventional secretion

Recently, we established Jps1 as an alternative carrier for heterologous proteins using the production of synthetic nanobodies against SARS-CoV-2 as a test case. We were successful in generating a functional bivalent nanobody directed against SARS-CoV-2 spike protein RBD (Sy^68/15^-Jps1) (Philipp et al. 2021). However, nanobody export mediated by Cts1 is also of great interest due to its natural ability in mediating both the export of heterologous proteins and chitin binding. The latter property is of potential high value with respect to protein purification and immobilization. Thus, to test the dual applicability of Cts1, we first screened different nanobody-Cts1 fusions for their expression, unconventional secretion and binding activity against SARS-CoV-2 RBD using Sy^68/15^-Jps1 as a benchmark (Fig. 1A). To this end four different strains were generated that produce anti-RBD nanobody versions fused to Cts1. These nanobody versions included the two synthetic nanobodies generated by Wagner et al. (2020) as single entities (Sy^15^-Cts1, Sy^68^-Cts1) and two llama-derived nanobodies VHH E and VHH V (here termed VHH^E^ and VHH^V^) generated by König et al. (2021) (VHH^E^-Cts1, VHH^V^-Cts1). In addition, a strain expressing a hetero bivalent version, pairing VHH^E^ with VHH^V^, was designed, since these were shown to display synergistic activity (König et al. 2021) (VHH^VE^-Cts1). Finally, a strain for production of a double mono bivalent VHH^E^ version was generated to test the binding capability of dimers with identical antigen binding sites (VHH^EE^-Cts1). The published synthetic nanobody versions Sy^68/15^-Cts1 (no antigen binding activity) and bivalent Sy^68/15^-Jps1 (alternative carrier; shows binding activity), pairing two synthetic nanobodies with different antigen binding sites, served as controls (Philipp et al. 2021). *U. maydis* expression strains for all protein versions were generated in the background of laboratory strain AB33P8Δ lacking eight extracellular proteases to optimize secretory yield (Terfrüchte et al. 2018). Expression and secretion of all versions was investigated via Western blot analyses. In cell extracts fusion proteins of the expected sizes were present for all variants, however, huge differences in expression level were observed. The analysis of culture supernatants confirmed sufficient secretion for the variants Sy^15^-Cts1, VHH^E^-Cts1, VHH^V^-Cts1, VHH^EE^-Cts1 and Sy^68/15^-Jps1, again displaying strong variation in the detected amounts (Fig. 1 B, Suppl. Fig. 1 A, B). To test for RBD binding activity, cell extracts of all strains containing nanobody-Cts1 fusions were subjected to direct ELISA assays, using recombinant RBD as a bait and a commercial antibody sandwich for detection. The strongest binding was achieved for VHH^EE^-Cts1 and Sy^68/15^-Jps1 while VHH^E^-Cts1 showed about half the signal intensity. All other variants lacked clear volumetric binding activity (Fig. 1 C, Suppl. Fig. 1 C). Overall, significant binding activity could be demonstrated for 3 of the 8 nanobody variants and binding capabilities of nanobodies were improved in the multimerized variant of VHH^EE^-Cts1.

**Figure 1:**
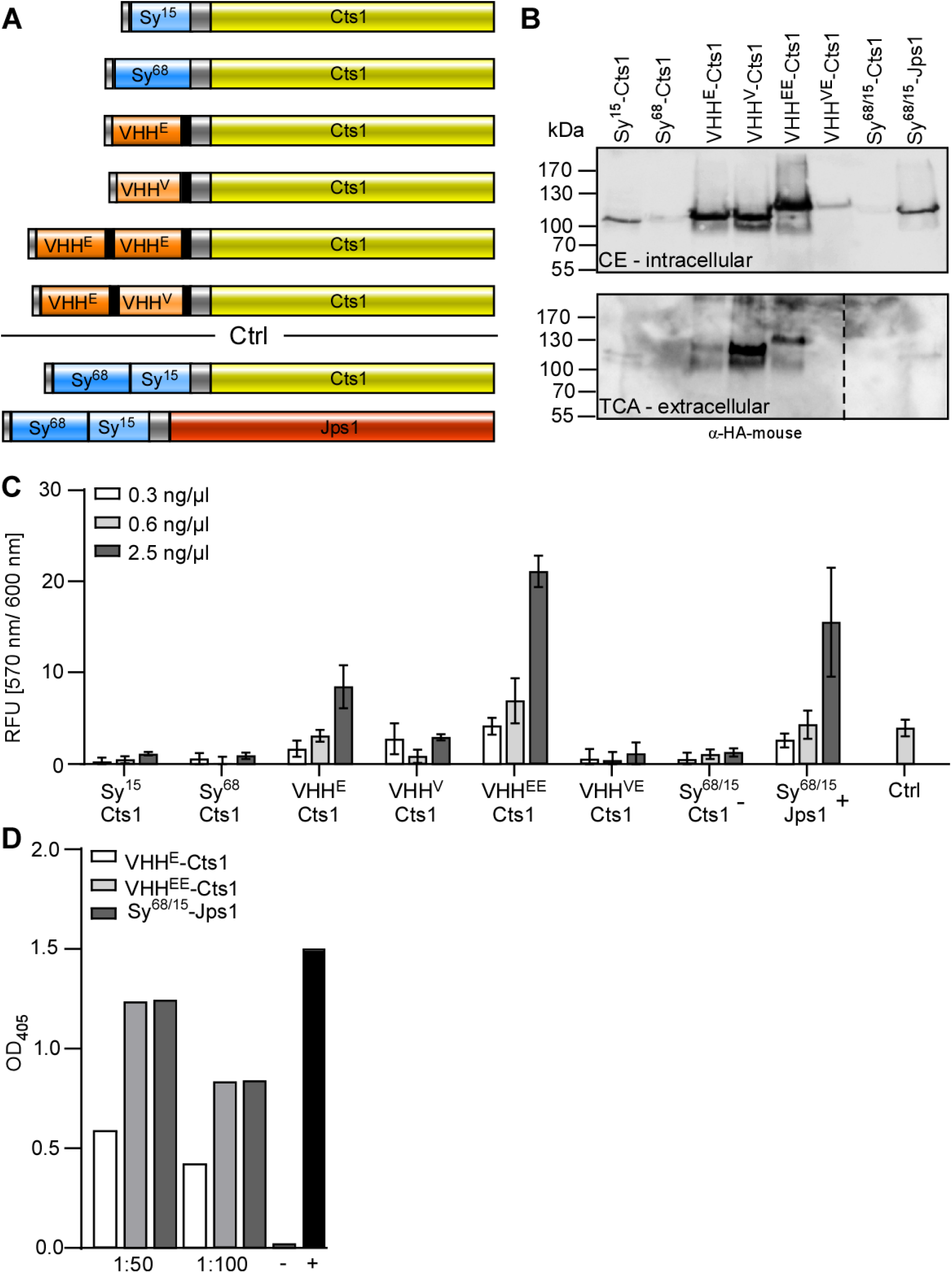
Functional screen of anti-SARS-CoV-2 nanobody-Cts1-fusion variants. **(A)** Schematic representation of nanobody protein variants fused to chitinase Cts1 as a carrier for unconventional secretion. Synthetic nanobodies Sy^15^ and Sy^68^ as well as llama-derived nanobodies VHH^E^, VHH^V^ and bivalent VHH^VE^ as well as a tandem VHH^E^ (VHH^EE^) were fused to Cts1 (yellow) via an HA-tag (grey) for detection. A His-tag (grey) was added at the N-terminus as a purification tag. In the case of the VHH^E^ and VHH^V^ nanobodies GS-linkers (black) introduced by (König et al. 2021) were placed between individual nanobodies and between the nanobodies and Cts1. Sy^68/15^-Cts1 and Sy^68/15^-Jps1 (Philipp et al., 2021) dealt as negative and positive control, respectively. Protein schemes drawn to scale. **(B)** Western blot analysis to detect nanobody expression and secretion levels. Top, cell extracts: 10 μg of cell extracts producing the indicated protein variants were subjected to Western blot analysis. Bottom, culture supernatants of strains producing indicated protein variants. Proteins were enriched from the supernatant via TCA precipitation, the HA tag was used for detection. Nanobody-Cts1-fusions were detected using an HA-mouse antibody and migrate slightly above their expected sizes around 100 kDa. **(C)** Direct ELISA of nanobody-Cts1-fusions against 1 μg/well of RBD domain coated to ELISA plate and detected by a sandwich of anti-HA (mouse) and an anti-mouse-HRP conjugate. Cell extracts of indicated expression strains were added to wells in serial dilutions of 0.3 ng/μl, 0.6 ng/μl and 2.5 ng/μl. The experiment was carried out in three biological replicates. Error bars depict standard deviation. **(D)** Direct ELISA of nanobody-Cts1 fusions against full length spike protein directly detected with an anti-HA-HRP conjugate. Purified nanobody-fusions (100 μg/ml) were added to wells in dilutions of 1:50 and 1:100 in technical triplicates. One biological replicate is shown. (+) and (-) indicate controls included in QuantiVac ELISA-kit.

To further substantiate the results, the three most promising nanobody versions VHH^EE^-Cts1, VHH^E^-Cts1 and Sy^68/15^-Jps1 were purified and subjected to direct ELISA against the full length S1 protein using an Anti-SARS-CoV-2 QuantiVac ELISA-kit including controls (Fig. 1 D). All variants depicted significant binding activity with VHH^E^-Cts1 binding at concentrations of 5 ng/μl and 10 ng/μl and both VHH^EE^-Cts1 and Sy^68/15^-Jps1 showing more than two-fold elevated binding compared to VHH^E^-Cts1. In essence, two functional nanobody-Cts1-fusion proteins were obtained which were comparable to the current benchmark Sy^68/15^-Jps1, recognizing both recombinant RBD and full length S1 protein.

### 2.2 *In vivo* activity of nanobody-Cts1 fusions

To determine if *in vitro* binding to SARS-CoV-2 RBD translates to binding or even virus neutralization *in vivo*, adjusted neutralization assays were applied. These assays are widely used to test sera of vaccinated or recovered patients for SARS-CoV-2 neutralizing antibodies (Matusali et al. 2021, Müller et al. 2021). In this adjusted neutralization assay, infectious SARS-CoV-2 viral particles were diluted two-fold starting at 100 TCID50 (tissue culture infectious dose 50). Dilutions were then pre-incubated with the purified functional nanobody variants VHH^E^-Cts1, VHH^EE^-Cts1 and Sy^68/15^-Jps1. The mixtures were subsequently used to inoculate Vero cell cultures displaying the ACE2 receptor on their surface to analyze viral replication. To quantify infection and neutralization, qPCR analysis was carried out for each replicate at the onset of infection and at three days’ post infection. VHH^E^-Cts1 showed no virus neutralization with strongly declining Ct values between t0 and t3, indicating a replicative infection. VHH^EE^-Cts1 on the other hand showed neutralization up to 25 TCID50 and Sy^68/15^-Jps1 up to 50 TCID50 as indicated by stable Ct values (Fig. 2 A). These results confirm the functionality of the nanobody fusions VHH^EE^-Cts1 and Sy^68/15^-Jps1 even towards infectious virus. Given that binding does not necessarily reflect neutralization, but neutralization definitely includes binding, this also confirms that these two versions are capable of binding SARS-CoV-2 *in vivo*.

**Figure 2:**
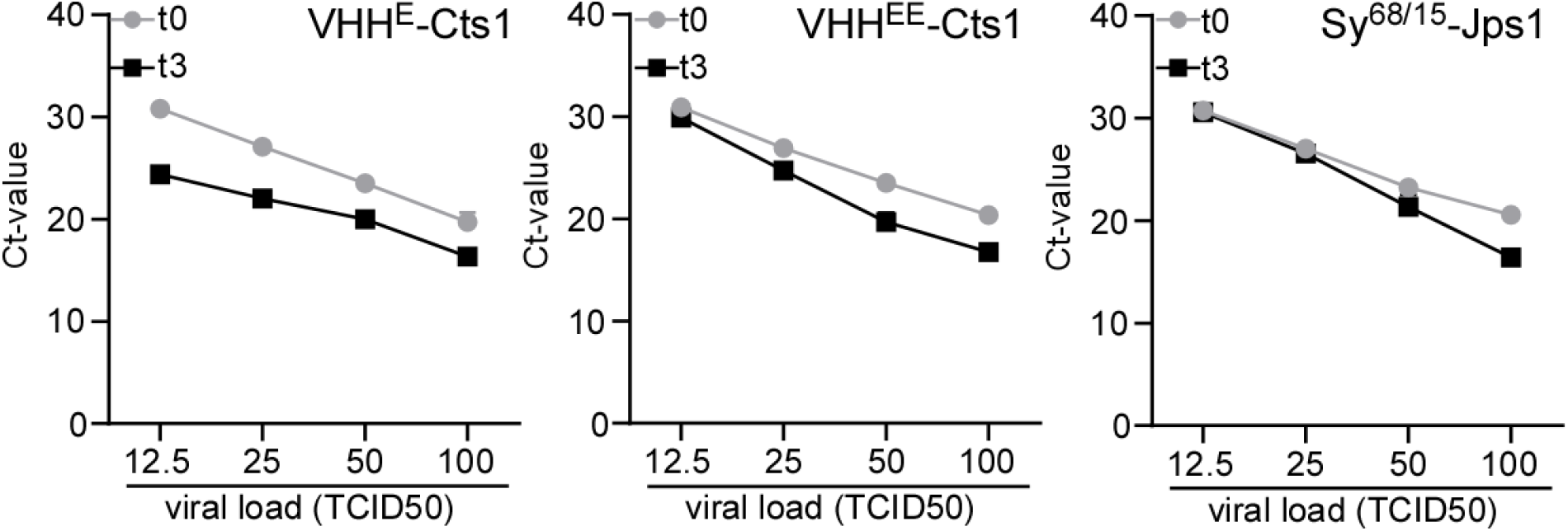
Neutralization assays conducted with anti-SARS-CoV-2 nanobody fusions produced in *U. maydis*. **(B)** qPCR analysis of infected mammalian cells to detect viral RNA in cell cultures treated with purified nanobody fusions/SARS-CoV-2 mixtures. Ct-values of samples at the onset of infection (t0) and three days’ post infection (t3) are depicted for each viral load and the respective applied nanobody fusion. Strong differences between Ct values of t0 and t3 indicate infection. Mean values of three biological replicates are depicted.

### 2.3 Characterization of Cts1 chitin binding and immobilization

Cts1 is capable of binding to chitin-coated surfaces like chitin magnetic beads without obvious degradation of the polymer (Terfrüchte et al. 2017). This observation could be developed into a strategy for a novel antigen test using an inexpensive surface based on bulk chitin obtained from crab shell or insects for immobilization of Cts1-nanobody fusions. To test chitin immobilization, we first recapitulated chitin binding on chitin beads using purified recombinant Cts1 (Fig. 3 A). Therefore, beads were mixed with recombinant Cts1 produced in *Escherichia coli*. After thorough washing, Cts1 was eluted from the beads, indicating stable binding and confirming previous results (Terfrüchte et al. 2017). Analysis of the fractions indicated that a significant amount of the protein was lost in the flow-through, suggesting that binding efficiency could be further improved in the future (Fig. 3 B). Next, β-glucuronidase (Gus)-Cts1 (Stock et al. 2012) obtained from *U. maydis* was used to quantify previous results (Terfrüchte et al. 2017) in the native system and to further characterize the chitin binding capacity of fusion proteins. Gus-Jps1 (Reindl et al. 2020), which is not predicted to bind to chitin, was used as a negative control (Fig. 3 C). Chitin beads were coated with the respective Gus-fusion proteins purified from *U. maydis* while washing and elution procedures were kept consistent to experiments carried out with recombinant Cts1. Indeed, Gus-Cts1 bound to chitin beads while no binding was observed for Gus-Jps1, confirming the binding capability of N-terminal Cts1-fusion proteins (Fig. 3 D). Quantification of signal intensities of the different fractions obtained in both experiments indicated that about 44% of the recombinant Cts1 and 68% of the native Gus-Cts1-fusion protein was captured by the beads (Fig. 3 E). To finally assay if the fusion protein is functional after immobilization, Gus activity was determined on chitin beads previously incubated with raw cell extracts of the Gus-Cts1 expression strain. Activity could specifically be detected on beads incubated with Gus-Cts1 containing cell extracts while the controls showed only background activity (Fig. 3 F). In essence, functional Cts1-fusion proteins can be immobilized on chitin beads and immobilization can even be achieved directly from raw cell extracts.

**Figure 3:**
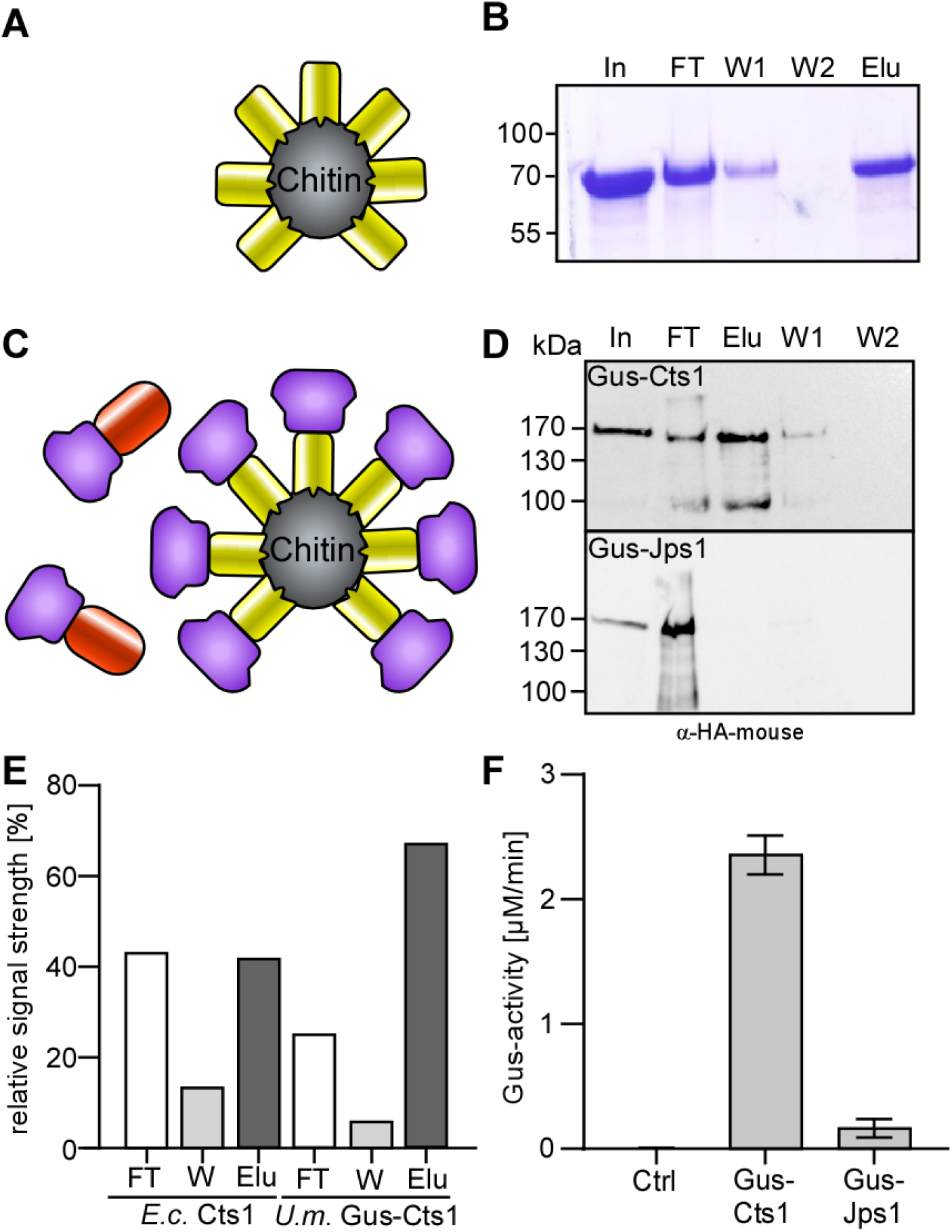
Chitin binding capabilities of Cts1 and Cts1-fusion proteins. **(A)** Experimental setup for initial Cts1 chitin binding experiments. *E. coli* derived, purified Cts1 (yellow) was coated to magnetic chitin beads, washed and subsequently eluted by boiling. **(B)** Coomassie-stained SDS-PAGE of different fractions obtained in the binding studies with recombinant Cts1 from *E. coli*: input (In), flow-through (FT), wash (W1 and W2) and elution (Elu) fractions of the experiment. **(C)** Experimental setup for Cts1 chitin binding experiments using *U. maydis* derived Gus-fusion proteins. Gus-Cts1 (purple-yellow) was coated to chitin beads, while a second set of beads treated with Gus-Jps1 (purple-red) dealt as a negative control. **(D)** Western blot analysis of input (In), flow-through (FT), wash (W1 and W2) and elution (Elu) fractions of purified Gus-Cts1 and Gus-Jps1-fusion protein incubated with chitin beads. **(E)** Relative quantification of SDS-PAGE and Western blot depicted in panels B and D. **(F)** On-bead Gus assays conducted with cell extracts of Gus-Cts1 and Gus-Jps1 incubated with chitin beads. After washing a Gus activity assay was conducted. Conversion from 4-MUG to 4-MU was monitored for 1 h. Cell extracts with Gus-Jps1 and of the progenitor strain (Ctrl) dealt as a negative control. Mean values of three biological replicates are shown. Error bars depict standard deviation.

### 2.4 Assessing the potential of anti-SARS-CoV-2 nanobody-Cts1 fusions for RBD capturing and chitin binding

To determine the capturing capabilities of the most promising nanobody variants VHH^E^-Cts1 and VHH^E^-Cts1, sandwich immunosorbent assays were conducted both on ELISA plates and non-classically on chitin beads. Sy^68/15^-Jps1 dealt as a control for both assays as it should show activity in plate-based ELISA but not on a chitin surface. In the first experiment, purified VHH^E^-Cts1, VHH^EE^-Cts1 and Sy^68/15^-Jps1 were coated to ELISA plates, incubated with serial dilutions of recombinant RBD and subsequently detected by a commercial RBD antibody and a cognate HRP conjugate (Fig. 4 A). In this plate-based ELISA all three nanobody variants were capable of capturing the RBD, however, only the most potent versions VHH^EE^-Cts1 and Sy^68/15^-Jps1 showed volumetric activity for serial RBD dilutions. As observed in the direct ELISA, VHH^EE^-Cts1 showed the strongest binding capability even at the lowest RBD concentration of 0.1 ng/μl after a detection time of 10 min, showing significantly stronger binding than VHH^E^-Cts1 which did neither reveal significant binding at 0.1 ng/μl nor volumetric activity with rising concentrations (Fig. 4 B). To determine if detection of RBD domain at similar concentrations can also be achieved using a chitin surface, chitin beads were incubated with purified VHH^E^-Cts1, VHH^EE^-Cts1 and Sy^68/15^-Jps1, mixed with RBD and again binding was detected using commercial antibodies (Fig. 4 C). Activity was obtained for both VHH^E^-Cts1 and VHH^EE^-Cts1, while no significant signal could be detected for Sy^68/15^-Jps1. As observed before, values for VHH^EE^-Cts1 were about doubled compared to those for VHH^E^-Cts1 (Fig. 4 D). In summary, these results demonstrate the potential of chitin-based ELISA using SARS-CoV-2 nanobody-Cts1-fusions and its specificity for the bifunctional Cts1 dealing as carrier and anchor for immobilization.

**Figure 4:**
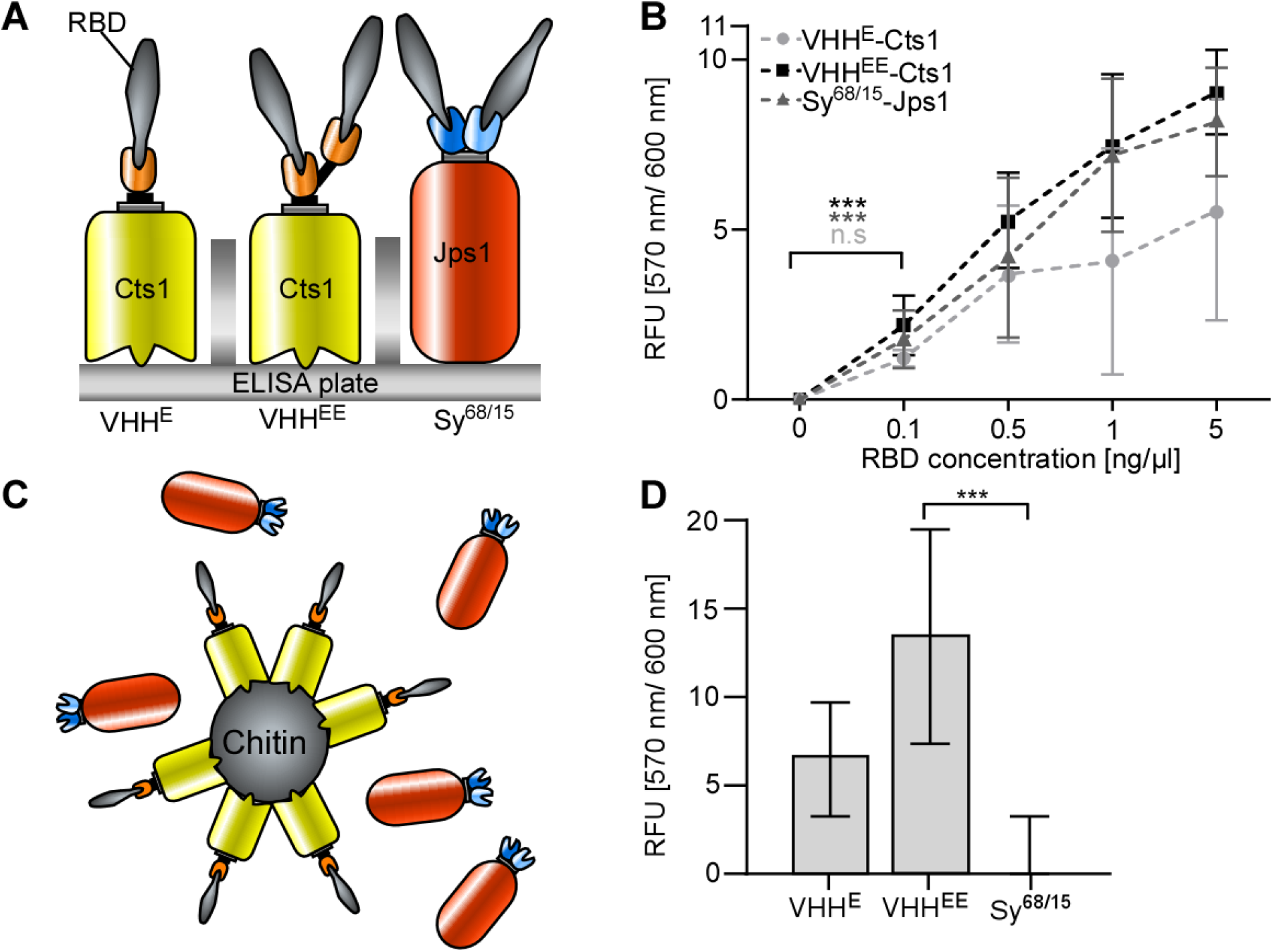
Plate-and chitin-based sandwich ELISA using Cts1-nanobody fusions for detection of SARS-CoV-2 RBD. **(A)** Experimental setup of plate-based sandwich ELISA. The indicated nanobody-Cts1 fusions were used as capture antibodies for serial dilutions of recombinant SARS-CoV-2 RBD (grey). **(B)** Quantitative results of plate-based sandwich ELISA. RBD (grey) was detected using an anti-RBD-(mouse) antibody and an anti-mouse-HRP conjugate. Mean values of three biological replicates are shown. Error bars depict standard deviation. Definition of statistical significance (***), p-value < 0.05. **(C)** Experimental setup of chitin-based sandwich ELISA test. The indicated nanobody-Cts1 fusions were coated to chitin beads to serve as capturing nanobodies, while Sy^68/15^-Jps1 dealt as negative control that is unable to bind to chitin. **(D)** Quantitative results of chitin-based sandwich ELISA. RBD was detected using an anti-RBD (mouse) antibody and anti-mouse-HRP conjugate. Sy^68/15^-Jps1 dealt as a negative control. Mean values of three biological replicates are shown. Error bars depict standard deviation. Definition of statistical significance (***), p-value < 0.05.

To further characterize the RBD capturing capabilities of the chitin-based detection system, the volumetric binding activity of the system was determined. Based on previous results, VHH^EE^-Cts1 was chosen as the most potent capturing nanobody. To this end, chitin beads were loaded with purified VHH^EE^-Cts1, subsequently incubated with recombinant RBD in serial dilutions and detected with a commercial antibody sandwich as described above (Fig. 5 A). A colorimetric reaction was obtained within a timeframe of two minutes, reflecting the rising input of the commercial RBD (Fig. 5 B).

**Figure 5:**
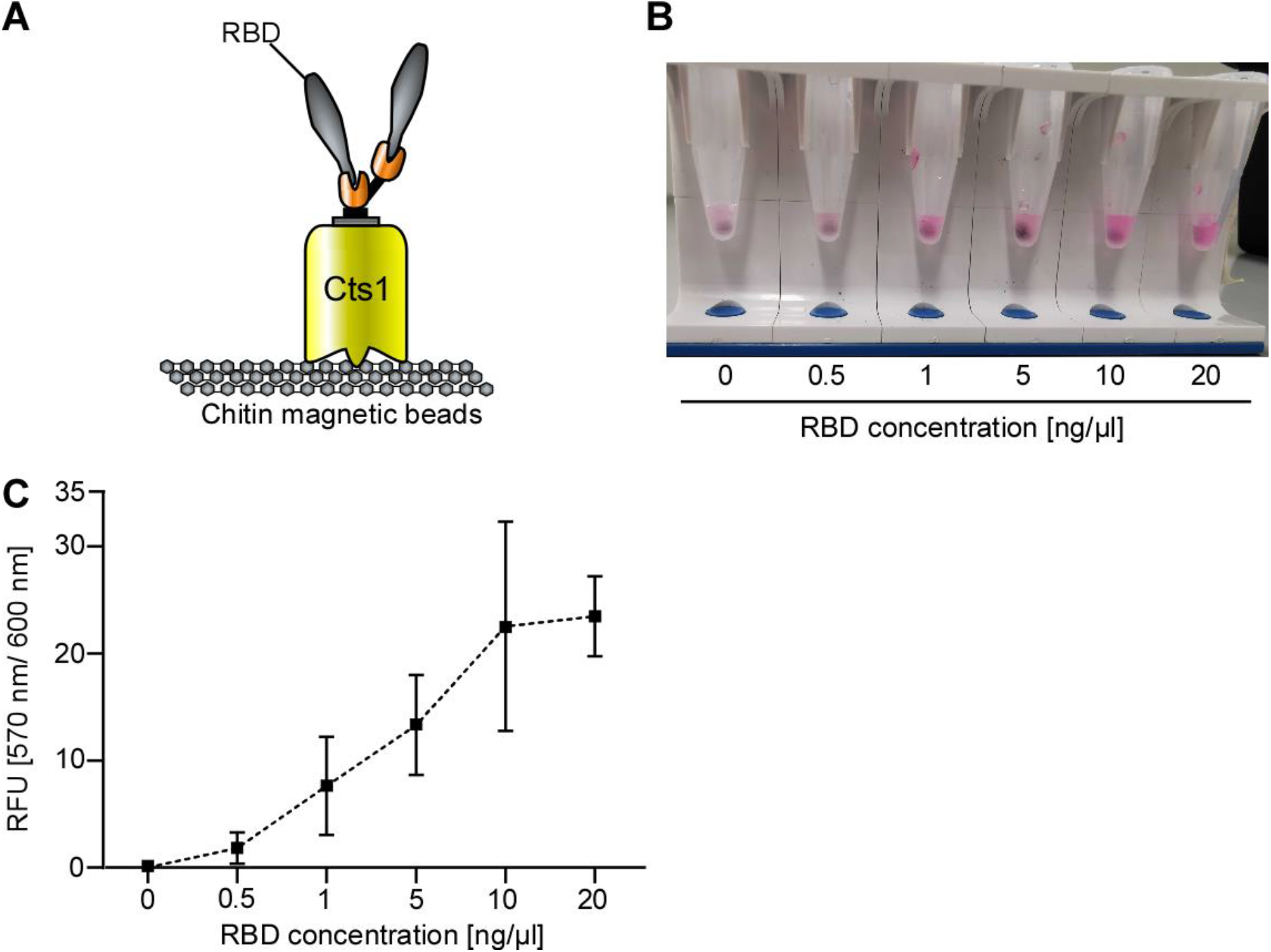
Chitin-based antigen test. **(A)** Setup of chitin-based ELISA. VHH^EE^-Cts1 binds to the chitin magnetic beads and is used as capture antibody for SARS-CoV-2 RBD. **(B)** RBD was added to VHH^EE^-Cts1 coated magnetic chitin beads in serial dilutions and subsequently detected using anti RBD (mouse) and anti-mouse-HRP antibodies. Picture depicts one representative replicate of the observed colorimetric reaction in reaction tubes. **(C)** Quantitative read-out of fluorescence measurements from chitin beads coated with VHH^EE^-Cts1, treated with serial dilutions of SARS-CoV-2 RBD. Mean values of three biological replicates are shown. Error bars depict standard deviation.

Quantitative fluorescence measurements of the samples confirmed these visual results (Fig. 5 C). Hence, overall, volumetric detection of SARS-CoV-2 RBD on a chitin surface is feasible, sensitive in the nanomolar range and in the given setup even faster than on a conventional ELISA plate.

## 3 Discussion

The aim of this study was to expand the repertoire of pharmaceutically relevant proteins produced in *U. maydis* towards versatile nanobodies for virus detection. To this end, Cts1-mediated secretion of active camelid-derived single- and bivalent nanobodies directed against the SARS-CoV-2 RBD was achieved. As an important strategical step towards industrial application, the produced nanobody fusions were immobilized on chitin, exploiting the natural capabilities of Cts1 and thus enabling detection of the cognate antigen via chitin-based ELISA.

The utilization of protein tags like Cts1 to export “passenger” proteins or peptides is a routine strategy in both bacterial and fungal hosts (Fleissner and Dersch 2010). Classical carriers are extracellular proteins that are naturally secreted in very high amounts and can be used as export proteins to guarantee high yields of secreted fusion protein. One such example are carbohydrate-active proteins like CBH1 from *Trichoderma reesei* (Zhong et al. 2011). As a positive side-effect, these protein tags can even enhance the solubility like shown for carbohydrate-binding module 66 (CBM66) in *E. coli* (Ko et al. 2021). In our study, different combinations of nanobody and Cts1 carrier resulted in varying levels of expression, secretion and activity. In addition, we observed a strong impact of the carrier choice on activity of the fused nanobody, confirming our earlier study (Philipp et al. 2021). While this underlines the necessity of a screening step of several constructs for each nanobody, it is in line with results obtained in other carrier based secretion systems (Wang et al. 2020b).

Similarly, also multiple powerful tags for protein purification and immobilization exist with the polyhistidin, the haemagglutinin or the SUMO tag as a few important examples (Porath et al. 1975, Field et al. 1988, Guerrero et al. 2015). The IMPACT system (New England Biolabs) even exploits a chitin-binding protein tag derived from *Mycobacterium xenopi* GyrA for protein purification with the intein tag (Chong et al. 1997, Chong et al. 1998). Importantly, in our system, chitinase Cts1 mediates both export and immobilization of the heterologous proteins. Thus, while normally carriers and tags for purification or immobilization are separated, Cts1 intrinsically combines both properties. This unique strategy will enable a very streamlined process design in the future.

Currently, we are using affinity chromatography to purify the Cts1-fusion proteins. In the future, the development of *in-situ* purification strategies for Cts1-fusion proteins from culture supernatant could greatly ease the purification process and thereby lower production costs of biopharmaceuticals. Similar non chromatographic purification processes have already proven successful using GST, biotin and streptavidin coated magnetic particles to purify protein from *E. coli cell* lysates (Franzreb et al. 2006) and supernatants (Fernandes et al. 2016) but also from human serum plasma (Santos et al. 2020).

In previous studies we had shown that chitin-coated beads are applicable for the purification of Cts1-fusion proteins (Terfrüchte et al. 2017). Now we expand on that and developed this interaction for nanobody immobilization in immunoassays. Since protein immobilization is generally achieved via protein adhesive polymers and not by specific protein-molecule interaction (Lin 2015, Andryukov 2020) this provides a novel tool towards inexpensive surface coating. The use of bio-based polymers for immobilization is of utmost interest since it allows for reduction of antigen test pricing and use of sustainable and inexpensive resources. To this end a similar study achieved SARS-CoV-2 detection based on nanobody immobilization on cellulose, albeit without using the immobilization tag for export at the same time (Sun et al. 2022).

Importantly, RBD capture capability of VHH^EE^-Cts1 in this study could be shown at concentrations in the low nanomolar range of 2.6 nm (0.1 ng/μl) in plate based ELISA, which is in the described range of other anti-RBD nanobodies between 0.9 nm and 30 nM (Weinstein et al. 2022), (Huo et al. 2020), (König et al. 2021). Moreover, published detection capacity of commercially available antigen tests is in the range of 0.65 pg/μl (nucleocapsid protein) to 5 ng/μl (spike protein) (Baker et al. 2020) (Grant et al. 2020). Thus, our chitin-based antigen detection system with a detection capacity of 0.5 ng/μl fits well into the described range, suggesting that it is competitive.

In a next step, we envision the application of our chitin immobilization strategy for nanobodies in virus detection for lateral flow assays. To date, detection of lateral flow assays is mostly enabled by colloidal gold particles (Oldenburg et al. 1998, Billingsley et al. 2017). Current investigation on chitin as a building block for nanocrystals and hydrogels (Xu et al. 2020, Gu et al. 2021), as well as initial experiments on drug loaded chitin scaffolds (Kovalchuk et al. 2019) demonstrates that generation of colloidal chitin particles loaded with nanobodies is a future possibility to further lower antigen test prices by exchanging gold as basic resource for detection by chitin.

Of note, we did not only verify the applicability of the nanobodies in virus detection but also successfully tested the neutralization of SARS-CoV-2 *in vivo*. This confirms nanobody binding of the infectious virus as opposed to only the RBD domain *in vitro*, which is necessary for antigen test application. The neutralizing activity could further motivate research towards drug development using unconventionally secreted proteins from *U. maydis* which as maize pathogen induces the formation of edible tumors and can thus be considered innocuous for humans (Juarez-Montiel et al. 2011). Nanobodies are currently discussed as novel drug targets due to ease of production, multimerization and favorable *in vivo* attributes, such as improved tissue penetration and decreased immunogenicity (Bannas et al. 2017, Salvador et al. 2019). Neutralizing mAbs are normally employed in biopharmaceutical cocktails in patients (Marrocco et al. 2019, Sun and Ho 2020). This strategy is applicable to nanobodies as well, however a study has shown comparable SARS-CoV-2 neutralization between a nanobody cocktail and a bivalent version of the same nanobodies in hamster models (Pymm et al. 2021), demonstrating that cocktails are not required, when bivalent constructs are used. *U. maydis* might be especially suited for the generation and production of larger multivalent constructs, given its ability to secrete huge proteins via the unconventional secretion route with a lack of *N*-glycosylation (Stock et al. 2012). Especially, since *N*-glycosylation has been shown to negatively affect pharmacokinetics of mAbs and even increased cytotoxicity before (Mastrangeli et al. 2020).

In summary, we provide a solid proof-of-principle for a chitin-based antigen test facilitated by components derived from unconventional secretion in *U. maydis*. We envision that in combination with sophisticated process engineering this technique could be developed into a lab-on-a-chip strategy (Zhuang et al. 2020). Thus, protein-based immobilization of nanobodies for target capture and detection are promising tools to develop alternative versatile and affordable technology for antigen testing.

## 4 Materials and methods

### 4.1 Molecular biology methods

All plasmids (pUMa/pUx vectors) generated in this study were obtained using standard molecular biology methods established for *U. maydis* including restriction ligation and Gibson cloning (Gibson et al. 2009). Enzymes for cloning were purchased from NEB (Ipswich, MA, USA). For the generation of pUMa4678 and 4679 *αgfpnb* was excised from pUMa2240 (Terfrüchte et al. 2017) by hydrolyzation with BamHI and SpeI. DNA sequences encoding for Sy^15^ and Sy^68^ (Walter et al. 2020) were amplified from synthetic gene blocks (IDT Coralville, IA, USA) using oligonucleotide pairs oAB908/oAB909 and oAB910/oAB911, respectively (Table 1). Subsequently PCR products were hydrolyzed with BamHI and SpeI and inserted into the backbone of pUMa2113 via restriction ligation cloning to generate pUMa4678 and 4679. Generation of pUx4 and pUx5 was achieved by excision of *αgfpnb* from pUMa2240 with BamHI and SpeI and amplification of *vhhe* and *vhhv* with BamHI and SpeI restriction sites from synthetic gene blocks using oligo nucleotide pairs oCD359/oCD360 and oCD363/oCD364, respectively. These sequences were subsequently hydrolyzed with BamHI and SpeI and inserted into the backbone of pUMa2240 via restriction ligation cloning, thereby generating pUx4 and pUx5. pUx6 was generated in a similar manner. However, after the hydrolyzation of the pUMa2240 backbone *vhhe* was amplified once with a BamHI and EcoRI and once with an EcoRI and SpeI hydrolyzation sites. After hydrolyzation two sequences for *vhhe* were inserted into the open reading frame via restriction ligation cloning, thereby encoding for fusion protein VHH^EE^-Cts1. For the generation of pUx7 this process was repeated but instead of using two *vhhe* sequences with differing hydrolyzation sites, the first *vhhe* sequence with BamHI and EcoRI hydrolyzation sites was exchanged for *vhhv* with corresponding hydrolyzation sites.

**Table 1.**
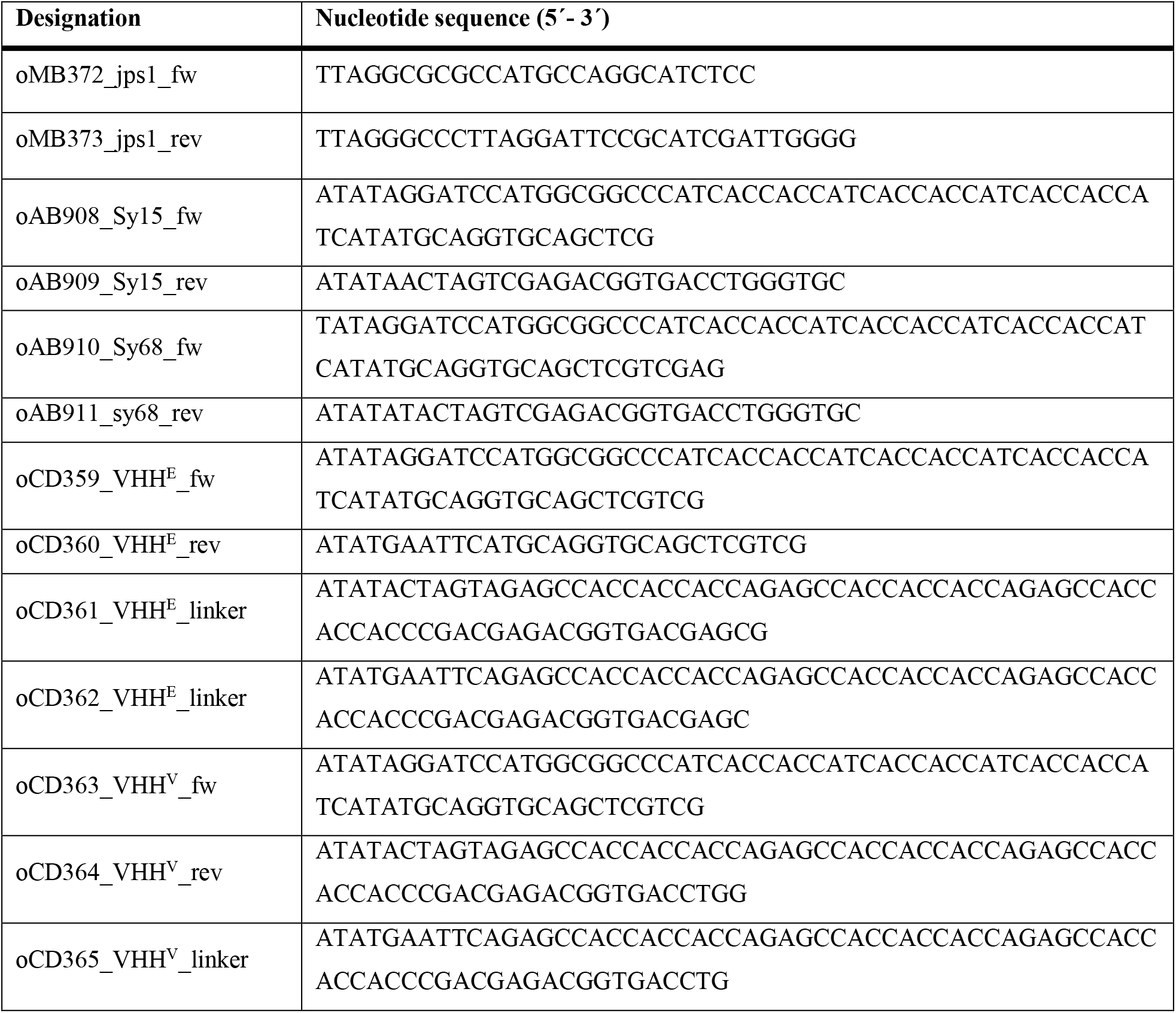
DNA oligonucleotides used in this study.

### 4.2 Strain generation

*U. maydis* strains used in this study were obtained by homologous recombination yielding genetically stable strains (Table 2). For genomic integrations at the *ip* locus, integrative plasmids were used (Stock et al. 2012). For genomic integration at the *ip* locus, integrative plasmids contained the *ip*^r^ allele, promoting carboxin (Cbx) resistance. Thus, plasmids were linearized within the *ip*^r^ allele using restriction enzymes SspI and SwaI to allow for homologous recombination with the *ip*^s^ locus. For all genetic manipulations, *U. maydis* protoplasts were transformed with linear DNA fragments. All strains were verified by Southern blot analysis. For *in locus* modifications the flanking regions were amplified as probes. For *ip* insertions, the probe was obtained by PCR using the primer combination oMF502/oMF503 and the template pUMa260. Primer sequences are listed in Table 1.

**Table 2.**
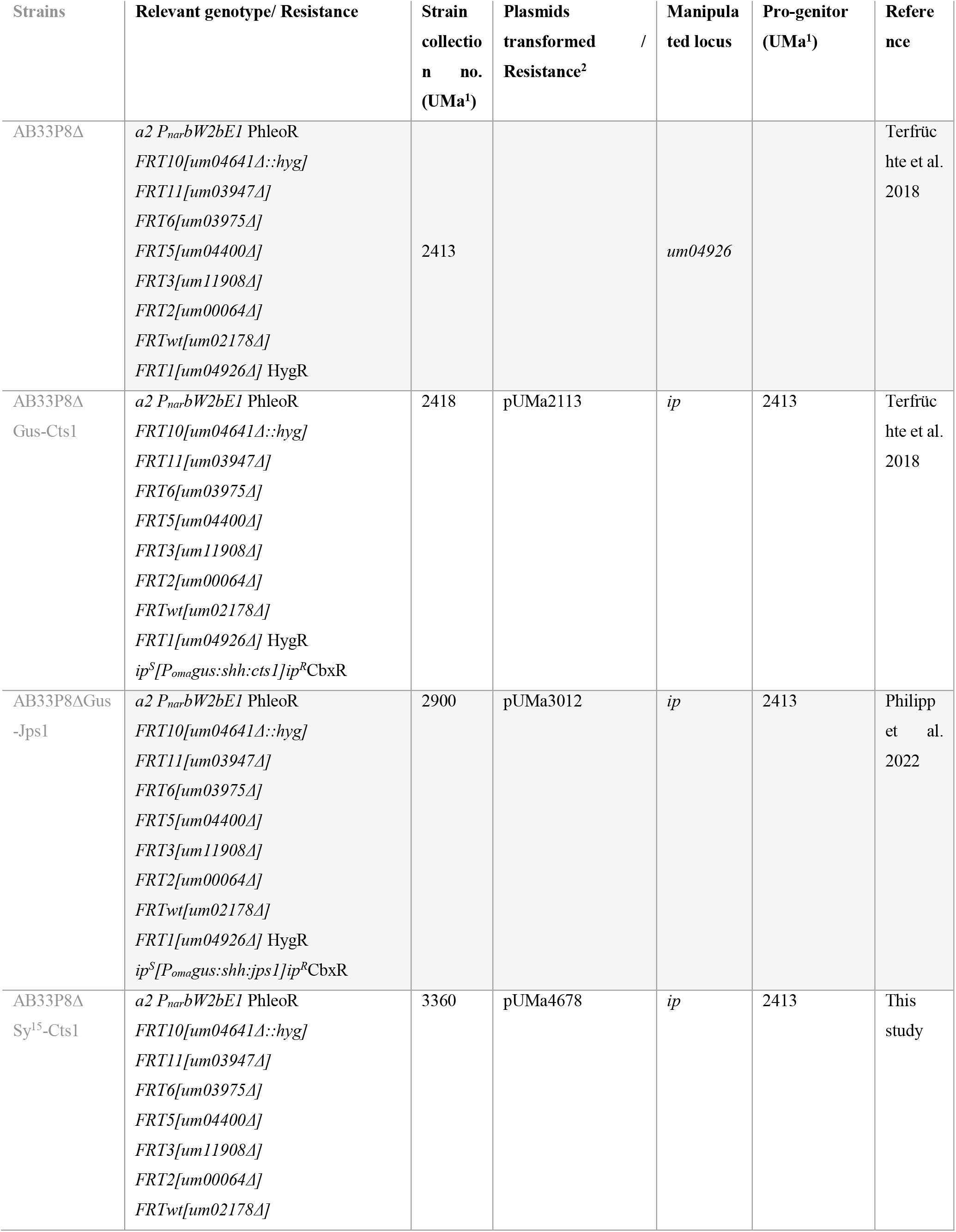
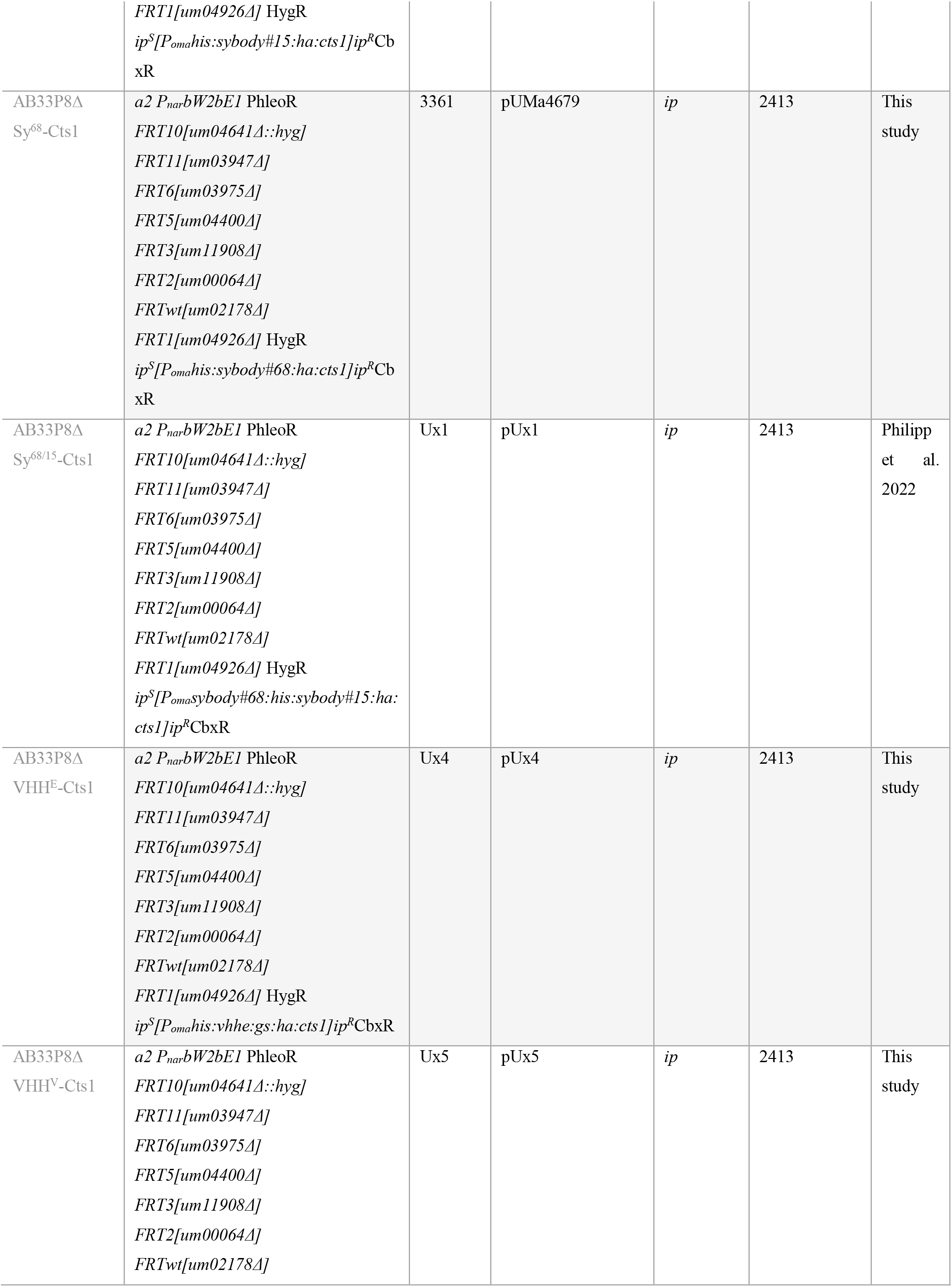
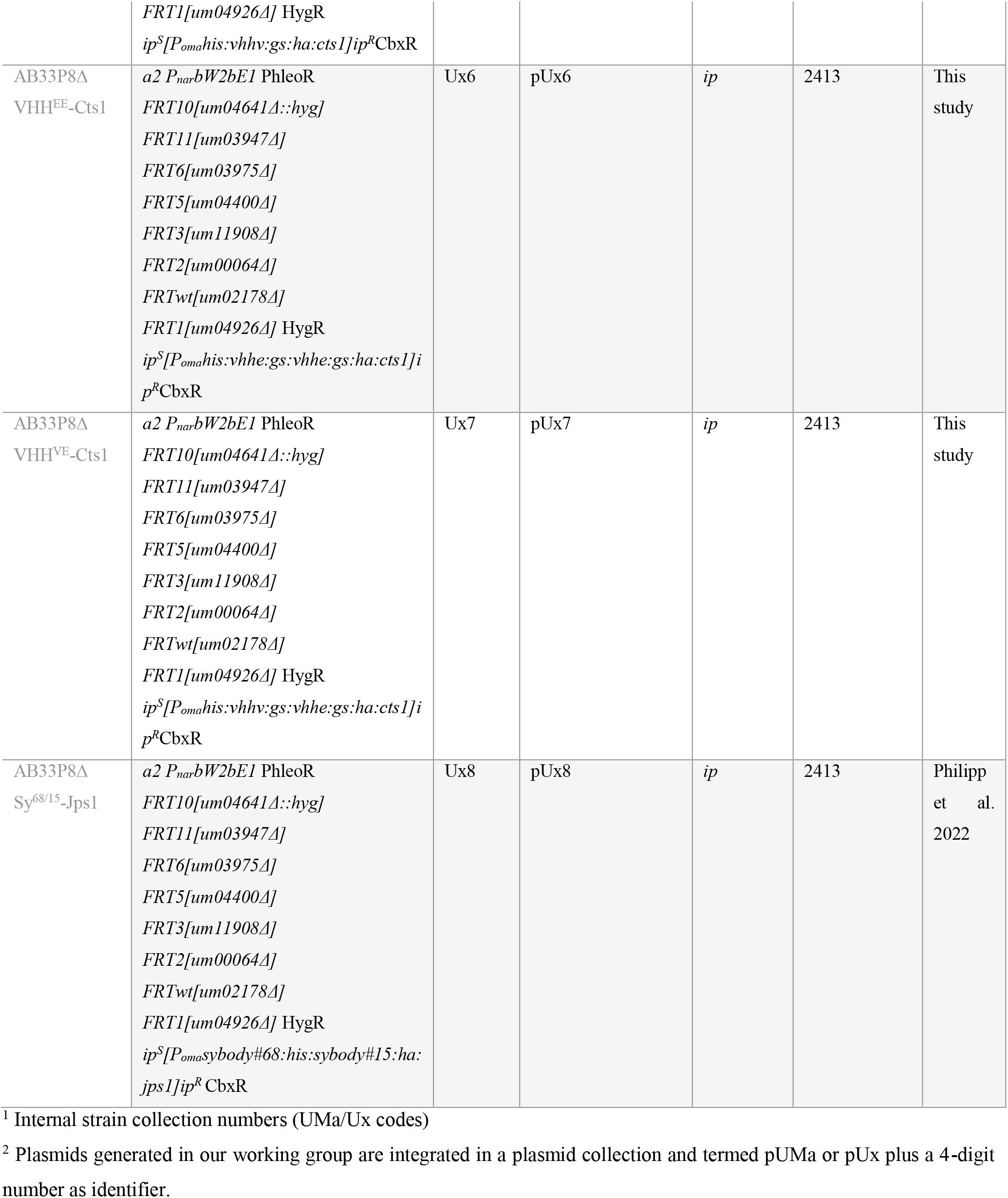
*U. maydis* strains used in this study.

### 4.3 Cultivation

*U. maydis* strains were grown at 28 °C in complete medium supplemented with 1% (w/v) glucose (CM-glc) if not described differently. Solid media were supplemented with 2% (w/v) agar agar. Growth phenotypes were evaluated using the BioLector microbioreactor (m2p-labs). MTP-R48-BOH round plates were inoculated with 1.5 ml culture per well and incubated at 1,000 rpm at 28 °C. Backscatter light with a gain of 25 or 20 was used to determine biomass.

### 4.4 Quantification of Gus activity on chitin beads

Gus activity was determined to quantify chitin binding of Gus-Cts1 using the specific substrate 4-methylumbelliferyl β-D galactopyranoside (MUG, Sigma–Aldrich). To his end 50 μg of *U. maydis* cell extracts were diluted in chitin binding buffer to a final volume of 500 μl. 50 μl chitin magnetic beads (New England Biolabs, Ipswich, MA, USA) were washed with 500 μl water, equilibrated with 500 μl chitin binding buffer (500 mM NaCl, 50 mM Tris-HCl buffer pH 8.0, 0,05 % Tween-20 (v/v)) and subsequently incubated with cell extracts in binding buffer at 4 °C on a stirring wheel for 16 h. Subsequently, chitin beads were washed with 500 μl chitin binding buffer and 500 μl of water, taken up in 2× Gus assay buffer (10 mM sodium phosphate buffer pH 7.0, 28 μM β-mercaptoethanol, 0.8 mM EDTA, 0.0042% (v/v) lauroyl-sarcosin, 0.004% (v/v) Triton X-100, 2 mM MUG, 0.2 mg/ml (w/v) BSA) and transferred to black 96-well plates. Relative fluorescence units (RFUs) were determined using a plate reader (Tecan, Männedorf, Switzerland) for 100 min at 28 °C with measurements every 5 minutes (excitation/emission wavelengths: 365/465 nm, gain 60). For quantification of conversion of MUG to the fluorescent product 4-methylumbelliferone (MU), a calibration curve was determined using 0, 1, 5, 10, 25, 50, 100, 200 μM MU.

### 4.5 Trichloroacetic acid precipitation

Gus-Cts1 and Gus-Jps1 secretion was analyzed by TCA precipitation of culture broths. Therefore, 2 ml of cultures grown in Verduyn medium (55.5 mM Glucose, 74.7 mM NH_4_Cl, 0.81 mM MgSO_4_×7H_2_O, 0.036 mM FeSO_4_×7H_2_O, 36.7 mM KH_2_PO_4_, 100 mM MES pH 6.5, 0.051 mM EDTA, 0.025 mM ZnSO_4_×7H_2_O, 0.041 mM CaCl_2_, 0.016 mM H_3_bBO_3_, 6.7 μM MnCl_2_×2H_2_O, 2.3 μM CoCl_2_×6H_2_O, 1.9 μM CuSO_4_×5H_2_O, 1.9 μM Na_2_MoO_4_×2H_2_O, 0.6 μM KI) to an OD_600_ of 3 were harvested by centrifugation at 11.000 × g and supernatant was transferred to a fresh reaction tube. 1 ml of cell free supernatants of cultures were chilled on ice, mixed with 400 μl 50% (v/v) TCA solution and incubated on ice at 4 °C overnight. Subsequently, protein pellets were harvested by centrifugation at 11.000 × g at 4 °C for 30 min. Supernatants were discarded and pellets were washed with 300 μl of −20 °C acetone followed by centrifugation at 11.000 × g at 4 °C for 20 min two times. Pellets were dried at room temperature and resuspended in Laemmli buffer containing 0.12 M NaOH. Resuspended pellets were denatured at 95 °C for 10 min and then subjected to SDS-PAGE and Western blot analysis.

### 4.6 Generation of cell extracts

For the verification of protein production via Western blot or further IMAC purification, cultures were grown to an OD600 of 1.0 and harvested at 5000 × g for 5 min in centrifugation tubes. Until further use, pellets were stored at −20 °C. For preparation of cell extracts, cell pellets were resuspended in 1 ml cell extract lysis buffer (100 mM sodium phosphate buffer pH 8.0, 10 mM Tris/HCl pH 8.0, 8 M urea, 1 mM DTT, 1 mM PMSF, 2.5 mM benzamidine, 1 mM pepstatin A, 2× complete protease inhibitor cocktail (Roche, Sigma/Aldrich, Billerica, MA, United States) and cells were crushed by agitation with glass beads at 2,500 rpm for 12 min at 4 °C. After centrifugation (11,000 × g for 30 min at 4°C), the supernatant was separated from cell debris and was transferred to a fresh reaction tube. For direct use protein concentration was determined by Bradford assay (BioRad, Hercules, CA, United States) (Bradford 1976). Otherwise, cell extracts were subjected to IMAC purification.

### 4.7 SDS PAGE and Western blot analysis

To assay protein production and secretion, 10 μg of cell extract or TCA precipitated samples were subjected to SDS-PAGE. SDS-PAGE was conducted using 10% (w/v) acrylamide gels. Subsequently, proteins were transferred to methanol activated PVDF membranes using semi-dry Western blotting. Nanobody fusion proteins were detected using a primary anti-HA (mouse; 1:3,000, Sigma-Aldrich, St. Louis, MO, USA). An anti-mouse IgG-horseradish peroxidase (HRP) conjugate (1:3,000 Promega, Fitchburg, United States) was used as secondary antibody. HRP activity was detected using the Amersham ^™^ ECL ^™^ Prime Western Blotting Detection Reagent (GE Healthcare, Chalfont St Giles, United Kingdom) and a LAS4000 chemiluminescence imager (GE Healthcare Life Sciences, Freiburg, Germany).

### 4.8 IMAC purification of His-tagged protein

Purification of *U. maydis* derived nanobody fusion proteins was achieved by generation of cell extracts from 400 ml of *U. maydis* culture harvested at an OD_600_ of 1.0 and subsequent Nickel^2+^-NTA purification. Therefore, culture harvested at 5000 *×* g for 5 min was resuspended in 8 ml lysis buffer (10 mM imidazole, 50 mM NaH_2_PO_4_, 300 mM NaCl, pH 8.0), 1.6 ml glass beads were added to cell suspension and cells were crushed by agitation with glass beads at 2,500 rpm at 4 °C for 12 min. Subsequently, cell debris was removed by centrifugation at 11,000 × g at 4 °C for 30 min. Nickel^2+^-NTA matrix was settled in empty columns and after flow-through of ethanol, equilibrated with 10 column volumes of lysis buffer. Subsequently, matrix was dissolved in cleared cell extracts and the mixture was incubated on a stirring wheel at 4 °C for 1 h. Subsequently, flow-through was discarded and matrix was washed with 5 column volumes of washing buffer (20 mM imidazole 50 mM NaH_2_PO_4_, 300 mM NaCl, pH 8.0). Protein was eluted in two fractions of 2 ml each using elution buffer 1 (lysis buffer, 150 mM imidazole) and elution buffer 2 (lysis buffer, 250 mM imidazole). For application in ELISA elution fractions were pooled via Amicon Ultra-15 50k centrifugal filter units (Merck Millipore, Burlington, MA, USA). Elution buffer was chosen for the intended application (coating buffer for sandwich ELISA, chitin binding buffer for chitin ELISA, PBS-T for direct detection, see chapters 4.10-4.11 for buffer composition).

### 4.9 *In vivo* neutralization assays

Nanobodies were IMAC purified and stored at 4 °C prior to incubation with SARS-CoV-2. Nanobodies at concentration of 0.5 mg/ml in PBS-buffer were incubated with SARS-CoV-2 particles in serial dilutions for 1 h at 37 °C. Subsequently, Vero cells (ATCC-CCL-81) displaying ACE2 were inoculated with pre incubated samples. After three days of incubation visual microscopic analysis was conducted using an Eclipse TS100 (Nikon, Minato, Japan) to observe cytopathic effects and thus determine if infection had occurred. qPCR analysis was conducted using anti-SARS-CoV-2 primer pairs specific to the E-gene (Corman et al. 2020) and Lightmix Modular SARS and Wuhan CoV E-gene (Roche Lifescience, Basel, Switzerland) in an ABI 7500 Fast PCR cycler (PE applied biosystems, Waltham, MA, USA).

### 4.10 Direct ELISA

For detection of nanobody binding activity protein adsorbing 384-well microtiter plates (Nunc^®^ Maxisorp^™^, ThermoFisher Scientific, Waltham, MA, USA) were used. Wells were coated with 1 μg Gfp for anti-GfpNB or 1 μg commercially available SARS-CoV-2 Spike-RBD-domain protein for SARS-CoV-2 nanobody-Cts1-fusion proteins (Invitrogen, Waltham Massachusetts, USA). Recombinant Gfp was produced in *E. coli* and purified by Ni^2+^-chelate affinity chromatography as described earlier (Terfrüchte et al. 2017). 1 μg BSA per well dealt as negative control (NEB, Ipswich, MA, USA). Samples were applied in a final volume of 100 μl coating buffer (100 mM Tris-HCL pH 8, 150 mM NaCl, 1 mM EDTA) per well at room temperature for at least 16 h. Blocking was conducted for at least 4 h at room temperature with 5% (w/v) skimmed milk in coating buffer. Subsequently, 5% (w/v) skimmed milk in PBS (137 mM NaCl, 2.7 mM KCl, 10 mM Na_2_HPO_4_, 1.8 mM KH_2_PO_4_, pH 7.2) were added to defined protein amounts of nanobody fusion protein samples purified from culture supernatants or cell extracts via Ni^2+^-NTA gravity flow and respective controls. 100 μl of sample was added to wells coated with the cognate antigen and BSA. The plate was incubated with samples and controls over night at 4 °C. After 3× PBS-T (PBS supplemented with 0.05% (v/v) Tween-20, 100 μl per well) washing, a primary anti-HA antibody (mouse, Sigma-Aldrich, St. Louis, MO, USA) 1: 5,000 diluted in PBS supplemented with skimmed milk (5% w/v) was added (100 μl per well) and incubated for 2 h at room temperature. Then wells were washed again three times with PBS-T (100 μl per well) and incubated with a secondary mouse-HRP antibody (goat, Promega, Madison, WI, USA) (50 μl per well) for 1 h at room temperature (1: 5,000 in PBS supplemented with skimmed milk (5% w/v)). Subsequently, wells were washed three times with PBS-T and three times with PBS and incubated with Quanta Red™ enhanced chemifluorescent HRP substrate (50:50:1, 50 μl per well, ThermoFisher Scientific, Waltham, MA, USA) at room temperature for 15 min. The reaction was stopped with 10 μl Quanta Red^™^ stop solution per well and fluorescence readout was performed at 570 nm excitation and 600 nm emission using an Infinite M200 plate reader (Tecan, Männedorf, Switzerland).

For ELISA against the full-length spike protein, experiments were carried out with the Anti-SARS-CoV-2-QuantiVac-ELISA (IgG)-Kit (Euroimmun, Lübeck, Germany) according to the manual. Controls were detected using the secondary anti-human-HRP antibody delivered with the Kit. Nanobody fusions were detected using anti-HA-HRP (Miltenyi Biotec, Bergisch Gladbach, Germany).

### 4.11 Sandwich ELISA

To determine nanobody-Cts1-fusion capabilities to act as capture antibody for an antigen test application, a mixture of 0.5 μg of IMAC purified protein and 0.5 μg BSA (New England Biolabs, Ipswich, MA, USA) in 100 μl of coating buffer per well was added to 384-well microtiter plates (1 μg without BSA for direct detection). Coating was conducted for 16 h at 4 °C. Subsequently, plates were blocked with 5% skimmed milk in coating buffer for 2 h at room temperature. RBD samples were added in serial dilutions in a volume of 100 μl sample buffer (5% skimmed milk powder in PBS-T) and incubated for 2 h at room temperature. Subsequently plates were washed 3 times with PBS-T and primary antibody (anti-RBD-mouse, R&D systems, Minneapolis, MN, USA) was added in a dilution of 1: 5,000 in sample buffer and incubated for 2 h at room temperature. Afterwards wells were washed again with PBS-T thrice and incubated with secondary mouse-HRP antibody (goat, Promega, Fitchburg, WI, United States) was added in a dilution of 1: 5,000 in 50 μl sample buffer and incubated for 1 h at room temperature. Prior to detection plates were washed thrice with 100 μl PBS-T and three times with 100 μl PBS per well. Detection was carried out using Quanta Red^™^ enhanced chemifluorescent HRP substrate (50:50:1, 50 μl per well, ThermoFisher Scientific, Waltham, MA, USA) at room temperature for 10 min. The reaction was stopped with 10 μl Quanta Red^™^ stop solution per well and fluorescence readout was performed at 570 nm excitation and 600 nm emission using an Infinite M200 plate reader (Tecan, Männedorf, Switzerland).

### 4.12 Chitin based sandwich ELISA

For chitin-based sandwich ELISA 50 μl of chitin magnetic beads (New England Biolabs, Ipswich, MA, USA) were transferred into a 1.5 ml reaction tube, washed with 500 μl of water and equilibrated in 500 μl of chitin binding buffer (500 mM NaCl, 50 mM Tris-HCl buffer pH 8.0, 0,05% Tween-20 (v/v)). Subsequently 2 μg of IMAC purified protein was added in a final volume of 500 μl chitin binding buffer. Coating was conducted on a stirring wheel at 4 °C for 16 h. Afterwards chitin beads were blocked with 5% skimmed milk powder in chitin binding buffer on a stirring wheel at room temperature for 2 h. In the next step chitin beads were washed thrice with PBS-T, RBD samples were added in serial dilutions in a volume of 100 μl ELISA sample buffer and incubated on a stirring wheel at room temperature for 2 h. After removal of the sample buffer chitin magnetic beads were taken up in 100 μl PBS-T, transferred to a fresh reaction tube and subsequently washed three times with 500 μl PBS-T before addition of primary antibody (R&D systems, Minneapolis, MN, USA) 1:5000 in 200 μl sample buffer. The primary antibody was incubated with chitin magnetic beads on a stirring wheel at room temperature for 2 h. Subsequent to primary antibody removal chitin magnetic beads were washed three times with PBS-T and incubated with secondary mouse-HRP antibody (goat, Promega, Fitchburg, United States) 1:5000 in 100 μl sample buffer on a stirring wheel at room temperature for 1 h. For detection chitin magnetic beads were washed three times with 500 μl PBS-T and three times with 500 μl PBS before being taken up in 100 μl Quanta Red^™^ enhanced chemifluorescent HRP substrate (50:50:1, 50 μl per well, ThermoFisher Scientific, Waltham, MA, USA) and transferred to a black 96-well microtiter plate. Fluorescence readout was performed 2 min after addition of the substrate at 570 nm excitation and 600 nm emission using an Infinte M200 plate reader (Tecan, Männedorf, Switzerland) after stopping of the reaction with 10 μl QuantaRed^™^ stop solution.

## 5 Author contributions

M.P., L.M, M.A. and K.P.H designed the experiments. K.P.H. conducted initial chitin binding experiments (Fig. 3 B). L.M. conducted neutralization experiments and RT-PCR (Fig. 2). M.A. conducted direct ELISA against full length S1 (Fig. 1 D) M.P. purified nanobodies for neutralization experiments and direct S1-ELISA and conducted all other experiments. M.P. designed and prepared the figures and tables. K.S., H.S. and M.F. supervised the project. M.P. prepared the manuscript with advice from K.S.

## 7 Acknowledgements

We acknowledge B. Axler for excellent support in molecular cloning and strain generation.

